# Crowdsourced geometric morphometrics enable rapid large-scale collection and analysis of phenotypic data

**DOI:** 10.1101/023382

**Authors:** Jonathan Chang, Michael E. Alfaro

## Abstract

1. Advances in genomics and informatics have enabled the production of large phylogenetic trees. However, the ability to collect large phenotypic datasets has not kept pace.
2. Here, we present a method to quickly and accurately gather morphometric data using crowdsourced image-based landmarking.
3. We find that crowdsourced workers perform similarly to experienced morphologists on the same digitization tasks. We also demonstrate the speed and accuracy of our method on seven families of ray-finned fishes (Actinopterygii).
4. Crowdsourcing will enable the collection of morphological data across vast radiations of organisms, and can facilitate richer inference on the macroevolutionary processes that shape phenotypic diversity across the tree of life.

## Introduction

Integrating phenotypic data, such as anatomy, behavior, physiology, and other traits, with phylogenies is powerful strategy for investigating the patterns of biological evolution. Recent advances in next-generation sequencing (Meyer *et al.* 2008; Shendure & Ji 2008) and sequence capture technologies (Faircloth *et al.* 2012; Lemmon *et al.* 2012) have made phylogenetic inference of large radiations of organisms possible (McCormack *et al.* 2012, 2013; Faircloth *et al.* 2013, 2014). However, similar breakthroughs for generating new phenotypic datasets have been comparatively uncommon, likely due to the high expense and effort required (reviewed in Burleigh *et al.* 2013).

Creating these large phenotypic datasets has generally required an extended dedicated effort of measuring and describing morphological or behavioral traits that are then coded into a comprehensive data matrix. One such example is the Phenoscaping project (http://kb.phenoscape.org; Deans *et al.* 2015), and related efforts in the Vertebrate Taxonomy Ontogeny (Midford *et al.* 2013) and Hymenoptera Anatomy Ontology (Yoder *et al.* 2010), which require large amounts of researcher effort to collate. Other approaches include using machine learning (Dececchi *et al.* 2015), machine vision (Corney *et al.* 2012a; b), or natural language processing (Cui 2012) to identify or infer phenotypes. These statistical techniques function ideally with either a large training dataset (e.g., a predefined ontogeny database) or a complex model (Brill 2003; Halevy *et al.* 2009; Hastie *et al.* 2009), both of which also require intensive researcher effort to build and validate. Finally, methods such as high-throughput infrared imaging, mass spectrometry, and chromatography have been successfully used in plant physiology (Furbank & Tester 2011) and microbiology (Skelly *et al.* 2013), but these methods may not be applicable for zoological researchers. These approaches all share a similar goal of collecting large comparative datasets, but also require large investments in researcher effort. This bottleneck in researcher availability has limited the scope of work in comparative biology.

Although it is now possible to build phylogenetic trees with thousands of tips, and phenotypic data sets have similarly been growing larger and larger, the traits that are typically studied at this scale tend to be simple: geographic occurrences (Jetz *et al.* 2012), one or two continuous characters (Harmon *et al.* 2010; Rabosky *et al.* 2013), a single discrete character (Goldberg *et al.* 2010; Aliscioni *et al.* 2012; Price *et al.* 2012), or some combination of these (Pyron & Burbrink 2014; Zanne *et al.* 2014). A richer understanding of the forces that shape macroevolution requires the collection of more detailed phenotypic trait data at scale.

Here we present a method and toolkit to efficiently collect two-dimensional geometric morphometric phenotypic data at a high-throughput “phenomic” scale. We developed a novel web browser-based image landmarking application, and use Amazon Mechanical Turk (https://www.mturk.com) to distribute digitization tasks to remote workers (hereafter *turkers*) over the Internet, who are paid for their contributions. We evaluate the accuracy and precision of turkers by assigning identical image sets and digitization protocols to users who are experienced with fish morphology (hereafter *experts*), and compare the inter- and intra-observer differences between turkers and experts. To illustrate the efficiency of this approach, we construct a phylogenetic analysis pipeline to download photographs and phylogenies of seven actinopterygiian families from the web, collect Mechanical Turk shape results, analyze the rate of diversification and body shape evolution using BAMM (Rabosky 2014), and compare the time required for this workflow to traditional approaches. We also discuss the role that crowdsourcing is best suited in large-scale morphological analyses, and suggest ways to integrate crowdsourced data as part of larger initiatives to digitize biodiversity.

## Materials and methods

### Amazon Mechanical Turk

Amazon Mechanical Turk (“MTurk”) is a web-based service where Requesters can request work, known as Human Intelligence Tasks (“HITs”) to be performed by Workers. Workers work from home and submit the tasks over the Internet, where Requesters review it, and, if they are satisfied with the results, accept the work and pay the Worker. We use MTurk as a platform to distribute our geometric morphometric tasks and financially compensate the worker accordingly. Scientific collection of data over MTurk and similar services has generally been limited to the fields of psychology and computer science, and there have been few attempts to crowdsource biological trait data (Burleigh *et al.* 2013).

### Web-based geometric morphometrics

We developed an geometric morphometric digitization application that runs completely on the user’s local web browser, using the HTML5 Canvas interface. This simplifies the infrastructure challenge of needing to serve many crowdsourced workers simultaneously, since workers will not need to download desktop software such as tpsDig (http://life.bio.sunysb.edu/ee/rohlf/software.html) before generating data. The web application is configured with a simple JavaScript Object Notation (JSON) file that describes the landmarks necessary to complete an image digitization task (Supplemental Figure S1). Point landmarks, semilandmark curves, and linear measurements are all supported. The software is available at https://github.com/jonchang/eol-mturk-landmark.

Although digitizing and landmarking a single image (microtasks *sensu* Good & Su 2013) is effective for high-throughput work on MTurk, it is unsuitable for conducting controlled experiments. To solve this issue we also created a server-side application backend that automatically distributes tasks according to a configurable set of images and experimental protocol. This application mimics an official Amazon Mechanical Turk interface endpoint, to facilitate drop-in replacement for an existing MTurk workflow. External non-MTurk workers can also participate in the same experiment, ensuring consistent comparisons across separate groups. The software is available at https://github.com/jonchang/fake-mechanical-turk.

### Reliability analysis

Collecting landmark-based geometric morphometric data at scale permits detailed analysis of different sources of error, such as among- and within-observer variation (Von Cramon-Taubadel *et al.* 2007). To assess whether the quality of data gathered by workers recruited through Amazon Mechanical Turk was significantly different than traditionally-collected data, we asked turkers (*n* = 21) and experts (*n* = 8) to landmark a set of five fish images, five times each. All participants used the same protocol and same software to digitize the same set of fishes. The landmarks were carefully selected based on previously-published literature concerning fish shape (Supplemental Figure S2; Fink & Zelditch 1995; Cavalcanti *et al.* 1999; Rüber & Adams 2001; Klingenberg *et al.* 2003; Chakrabarty 2005; Frédérich *et al.* 2008; Claverie & Wainwright 2014; Thacker 2014). We also ensured that the chosen landmarks included morphological features that were relatively straightforward to digitize (the position of the eye) and features that were likely to be more challenging to digitize (the position of the preopercle bone), in order to test for turker and expert differences over a spectrum of difficulties. We report the inter-observer reliability for turkers and experts by computing the ratio of the among-individual and the sum of the among-individual and measurement error variance components in a repeated measures nested MANOVA (Palmer & Strobeck 1986; Zelditch *et al.* 2012).

To assess the differences between turker and experts on a per-landmark basis, we first compared the median turker position to the median expert position of each landmark. We assumed that the expert median was the true position of that landmark, and calculated the absolute Euclidian distance. Larger distances would indicate low turker accuracy, while smaller distances would indicate high turker accuracy. We then examined the variance in turker landmarks. For each landmark, we rotated the cloud of points to maximize variance in one dimension, and calculated the log-ratio of median absolute deviations (MAD) between turkers and experts. This rotation is a conservative approach for assessing the difference in variance between these two groups, because it maximizes any apparent differences in landmark position. A positive log-ratio indicated that experts had lower variance than turkers, while a negative log-ratio indicated that turkers had lower variance. For all subsequent analysis, we excluded landmarks where turkers performed especially poorly, where either the accuracy or precision components for a given landmark exceeded 1.5 times the interquartile range of that component.

To determine whether turkers and experts were statistically distinguishable, we performed a non-parametric MANOVA using the randomized residual permutation procedure (RRPP) with 1,000 iterations (Collyer *et al.* 2014). The RRPP method reduces the effect of the “curse of dimensionality” (*p >> n*, where the number of predictors greatly exceeds the number of observations), a common problem in geometric morphometrics, and has been shown to have increased statistical power compared to a method where the raw data are randomized instead (Anderson & Braak 2003). We test for a difference between mean turker and expert shapes against a null model of no difference between turker and expert changes, taking into account species-specific differences. A difference between models was considered significant if the p-value was less than *α* = 0.05.

As a separate test, we use linear discriminant analysis (LDA, Ripley 1996), a statistical classification algorithm that finds features to differentiate between different classes of data, in this case turkers and experts. We assessed the accuracy of the LDA classification using 10-fold cross validation (CV), which splits our data into 10 equally-sized groups, using nine for training and one for validation (Kohavi 1995; Hastie *et al.* 2009). An acceptable misclassification rate varies depends on application, but here we use a 25% misprediction rate as a standard for sufficient accuracy. This is a highly forgiving standard, since a 50% misprediction rate is no better than a coin flip, and a 25% misprediction rate would still erroneously classify one in four turkers as experts or vice versa. We also use quadratic discriminant analysis (QDA), which relaxes some of the assumptions of LDA, and similarly report the QDA misclassification rate.

We calculated the per-individual median shape for each species used, as well as the consensus turker and morphologist shapes, and projected these shapes into Procrustes space, to visualize the orthogonalized differences in median shape among and between the types of digitizers.

### Example: a phenomic pipeline for comparative phylogenetic analysis

A common strategy in fish comparative studies is to examine evolutionary dynamics within a single family (Ferry-Graham *et al.* 2001; Alfaro *et al.* 2005, 2007; Rocha *et al.* 2008; Hernandez *et al.* 2009; Dornburg *et al.* 2011; Frédérich *et al.* 2013; Santini *et al.* 2013; Sorenson *et al.* 2013; Claverie & Wainwright 2014; Thacker 2014), potentially due to the extensive amount of time necessary to collect data. To test whether our method can improve on the case where the data collection method is geometric morphometrics, we use the average time it took an expert to measure a single fish image and predict the time it would take for a single individual expert to measure all images at 5x replication, and compare it to the time it took turkers to collect these measurements at the same replication level. If the turkers in aggregate annotated images more quickly than a single expert would have, this suggests that the parallelization afforded by crowdsourcing is effective at reducing the total time required for data collection.

To demonstrate the utility of obtaining comparative data using this method, we use previously published phylogenies for seven fish families: Acanthuridae (Sorenson *et al.* 2013), Balistoidae, Tetraodontidae (Santini *et al.* 2013), Apogonidae, Chaetodontidae, Labridae (Cowman & Bellwood 2011; Choat *et al.* 2012), and Pomacentridae (Frédérich *et al.* 2013). We matched 147 species to left-lateral images from the Encyclopedia of Life (http://eol.org/) using their application programming interface (Parr *et al.* 2014). Crowdsourced workers placed landmarks describing body shape variation following a standard protocol (Supplementary Material). The Cartesian position of these landmarks were used in a generalized Procrustes analyses (Gower 1975; Rohlf & Slice 1990), which centers, scales, and rotates landmark configurations to minimize the least-squares distance between shapes. We then determined the major components of shape variation using a Procrustes-aligned principal components analysis (PCA) (Mardia *et al.* 1979; Bookstein 1991) with the R package *geomorph* (Adams & Otarola-Castillo 2013), and used these principal components axes for subsequent analyses.

We used Bayesian Analysis of Macroevolutionary Mixtures (BAMM; Rabosky 2014) to estimate rates of speciation and body shape evolution for all seven families. For the characters describing body shape, we use the PC axes whose eigenvalues exceeded the corresponding random broken-stick component (Jackson 1993; Legendre & Legendre 1998). BAMM estimates the location of rate shifts in either diversification or character evolution using a transdimensional (reversible jump) Markov Chain Monte Carlo method that samples a variety of models of lineage diversification and trait evolution. We assessed convergence and mixing using Tracer (Rambaut & Drummond 2007). We also repeated each analysis and simulated under the prior (without data) to exclude rate heterogeneity that occurred solely due to stochastic processes. We use a Bayes Factor criterion of *BF >* 5 to enumerate the set of credible shifts (Shi & Rabosky 2015) and visualized them in R using BAMMtools (Rabosky *et al.* 2015).

## Results

### Reliability analysis

For nearly all landmarks, turkers only differ from the expert consensus by a few tens of pixels (Figure 1, Supplemental Figure S3). The most accurate and precise points are those that are related to the position of the eye (landmarks E1 and E2). The least accurate are those in the opercular series (O1-O5), particularly the ones related to the preopercle (O1-O3) likely because in certain groups (e.g., Tetraodontidae) the preopercle is difficult to visualize from external morphology alone. Experts were generally more precise than turkers, however there were some landmarks where the turkers converged on very similar locations. Based on these results we exclude in subsequent analyses the landmarks relating to the distal margins of all fins (A3, A4, P3, P4, D3, D4), the preopercle bones (O1-O3), the dorsal fin for triggerfishes (D1, D2), and the opercular opening for pufferfishes (O4-O5), due to low turker accuracy.

**Figure 1:**
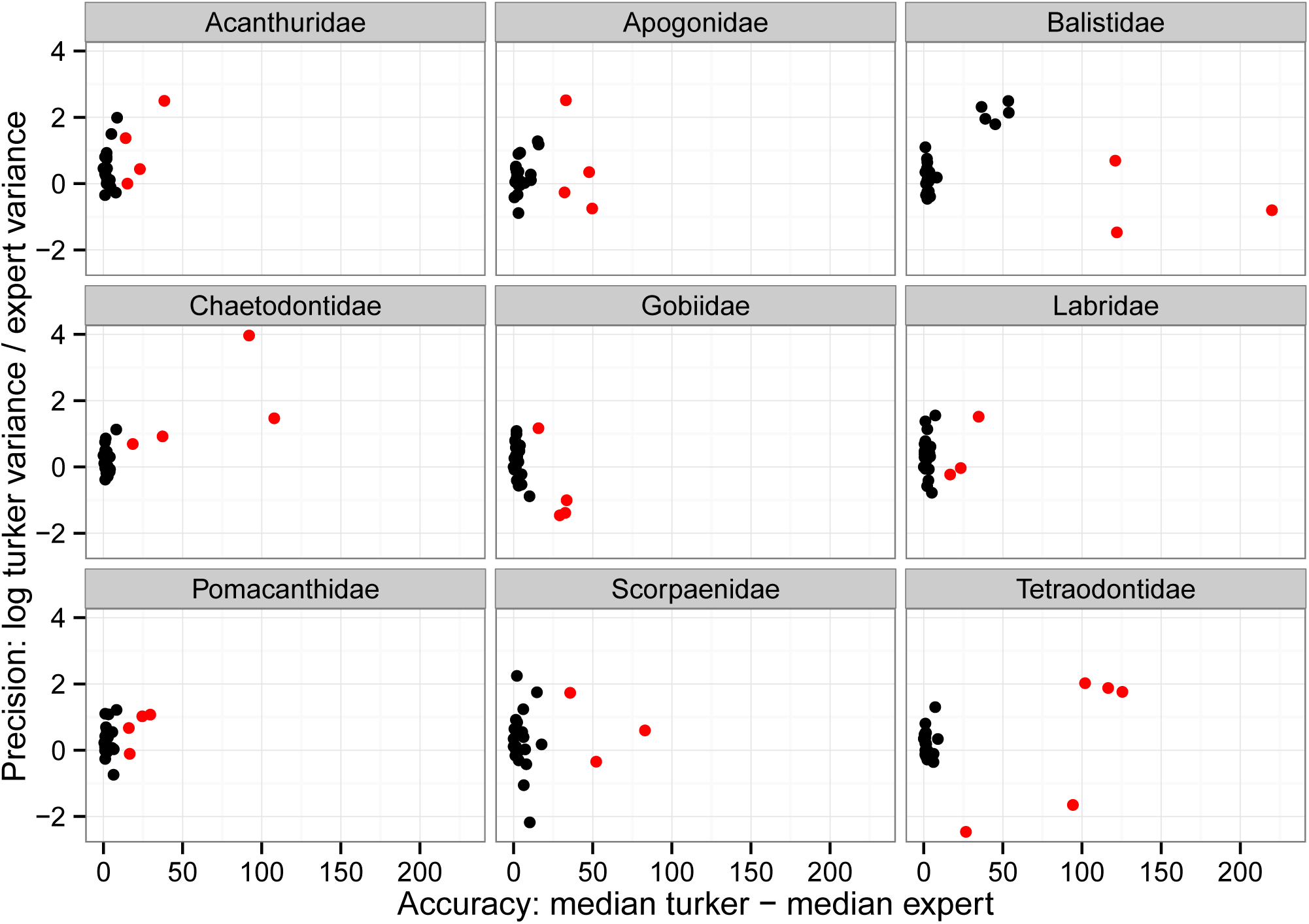
Per-family breakdown of accuracy vs. precision for each landmark. Accuracy is represented as the difference between the median turker location for that landmark and the median expert location, with the expert location assumed to be the true location. Precision is represented as the log-ratio of median absolute deviations between turkers and experts. More positive numbers indicate better expert precision, whereas more negative numbers indicate better turker precision. Points highlighted in red are those determined to be outliers (1.5 *imes* IQR). See Supplemental Information for a labeled version of this figure.

The inter-observer reliability of turkers and experts as measured by the ratio of among-individual and sum of the among-individual and measurement error ANOVA components was 96.4% and 90.9%, respectively. Although there is no current standard for acceptable levels of measurement reliability (Von Cramon-Taubadel *et al.* 2007), these percentages are not low enough to suggest pathologies in the measurement protocol.

The non-parametric MANOVA with RRPP failed to detect a significant difference between turker and expert shapes (*p* = 0.376, *Z* = 1.006007, *F* = 0.9938314). Similarly, both linear and quadratic distriminant analysis with 10-fold cross validation (Table 1) were unable to reliably distinguish between these two groups, for any given family. Although for some images the classifier showed slight improvement beyond a 50% coin flip, in all cases our model fell short based on a one in four (25%) acceptable misclassification rate. We conclude that, for any given sample of landmarks, it is challenging to statistically distinguish between expert-provided and turker-provided landmark configurations.

**Table 1:** Misprediction rate of linear discriminant analysis (LDA) and quadratic discriminant analysis (QDA) with 10-fold cross validation for each fish image. The discriminant model for each family was unable to meet the standard of one in four misclassifications, and in some cases, the more flexible QDA method performed worse than the LDA model.

| Family | LDA | QDA |
| --- | --- | --- |
| Acanthuridae | 0.446 | 0.370 |
| Apogonidae | 0.428 | 0.425 |
| Balistidae | 0.452 | 0.429 |
| Chaetodontidae | 0.438 | 0.424 |
| Gobiidae | 0.465 | 0.444 |
| Labridae | 0.416 | 0.382 |
| Pomacanthidae | 0.466 | 0.412 |
| Scorpaenidae | 0.496 | 0.468 |
| Tetraodontidae | 0.442 | 0.490 |

We projected turker and expert shape configurations into morphospace (Figure 2, Supplemental Figure S4) Although the overall space occupied by each family’s shape configurations vary, in practice, the aggregated median turker and expert shapes are not qualitatively different. The only exception is the triggerfishes (Balistidae), likely due to turker confusion over the exact location of dorsal fin due to their reduced anterior dorsal fin.

**Figure 2:**
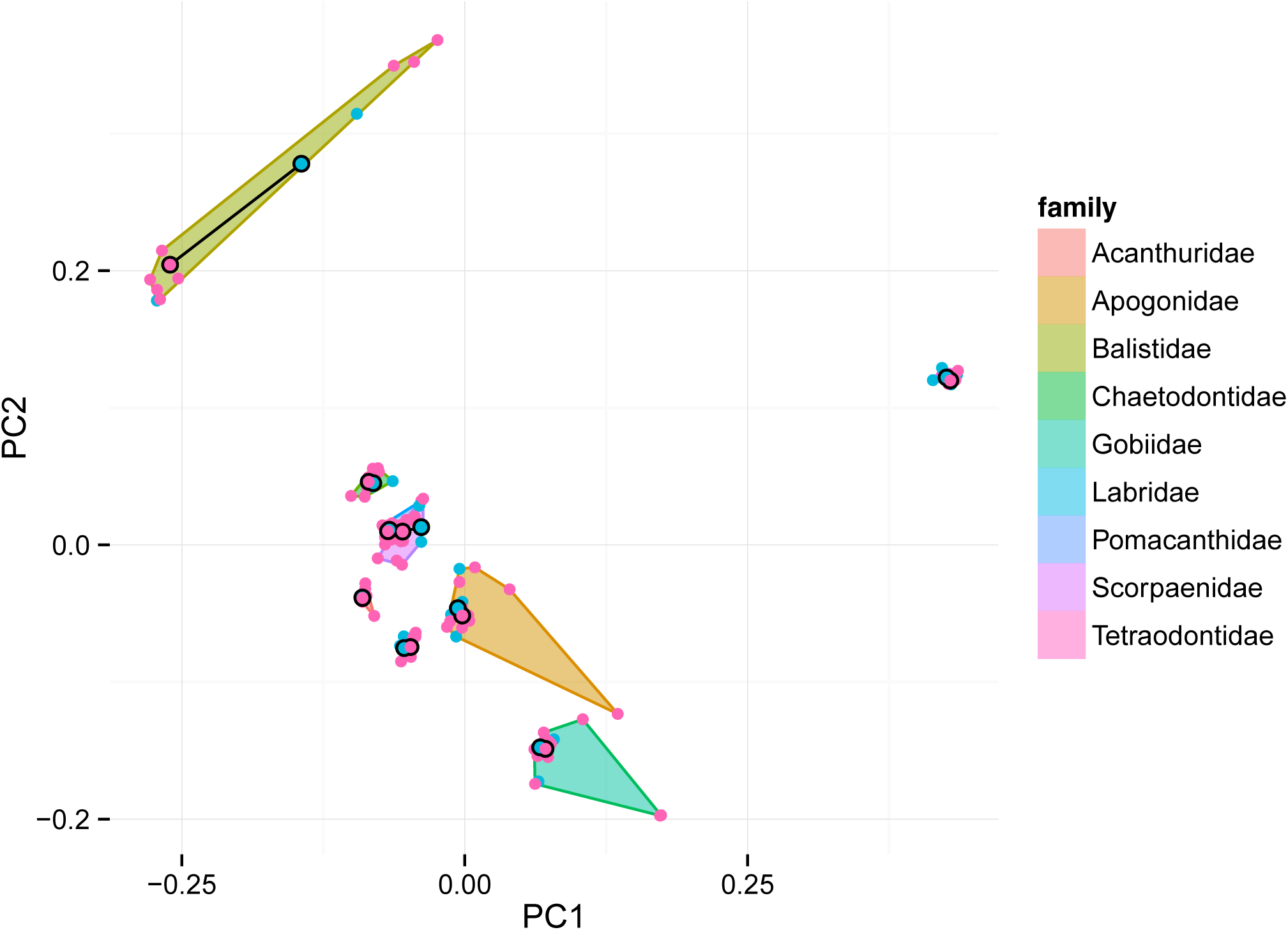
Morphospace projection for each observer’s mean shape. Blue points indicate experts, while red points indicate turkers. The mean shape for all turkers and experts for a given family is the point outlined in black for each family, and connected with a black line to help emphasize the difference between turker and expert mean shapes. The convex hull for each family is drawn to show the amount of among-observer shape variation.

**Figure 3:**
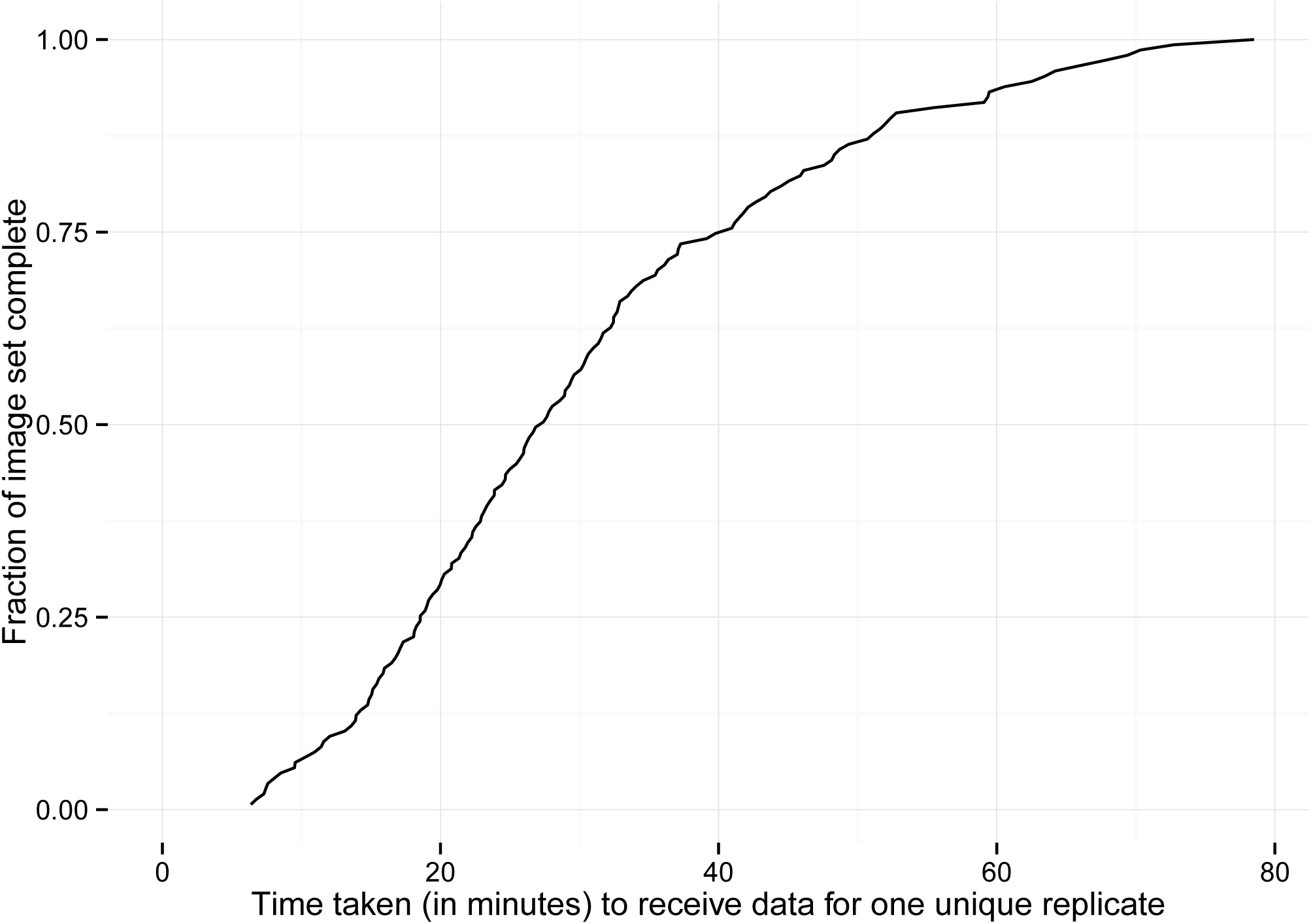
Line plot showing time to receive results for any given image (x axis) and the total fraction of the data set received (y axis). Landmarks were first received eight minutes after creation of the Amazon MTurk task, and at least one replicate was received for every image at the 80 minute mark.

### Phenomic pipeline for comparative phylogenetic analysis

Using a median expert time of 171.1s (∼2.85 minutes) per image, we estimate that a single morphologist would take 25151.7s (∼6.99 hours) to landmark all 147 images. At 5x replication, this would take 1772596s (∼20.52 days). By comparison, turkers took a total of 19789s (∼5.5 hours) to complete all images at 5x replication.

Using the broken-stick method of determining a PCA stopping point, we analyzed PC 1 through PC 5. We project per-species consensus shapes into Procrustes space (Figure 4, Supplemental Figure S5). The BAMMtools analysis uncovered substantial amounts of heterogeneity in the rate of body shape evolution and speciation in each family (Figure 5). Significant shifts in the rate of shape evolution or speciation were detected in three families: Labridae, Apogonidae, and Pomacentridae. The significant shifts in speciation rate corroborate those found in Cowman & Bellwood (2011) through either MEDUSA (Alfaro *et al.* 2009) or a relative cladogenesis statistic (Nee *et al.* 1992). Two significant shifts in shape evolution rate occur in the wrasses (Labridae). The first rate shift occurs deep in the tree, corresponding to the lineage containing the labrine, scarine, and cheiline tribes. The other shift is nested within that group, in *Sparisoma*. One shift in speciation rate also occurs in the wrasses, encompassing the genera *Chlorurus* and *Scarus*. One shift in speciation rate occurs in the cardinalfishes (Apogonidae), encompassing members of the genera *Apogon*, *Archamia*, *Zoramia*, *Ostorhinchus*, *Cheilodpterus*, *Gossamia*, *Fowleria*, and *Phaeoptyx* (Apogonini + Apogonichthynini *sensu* Mabuchi *et al.* 2014). One shift in the rate of shape evolution occurs in the damelfishes (Pomacentridae) in the genus *Amphiprion*.

**Figure 4:**
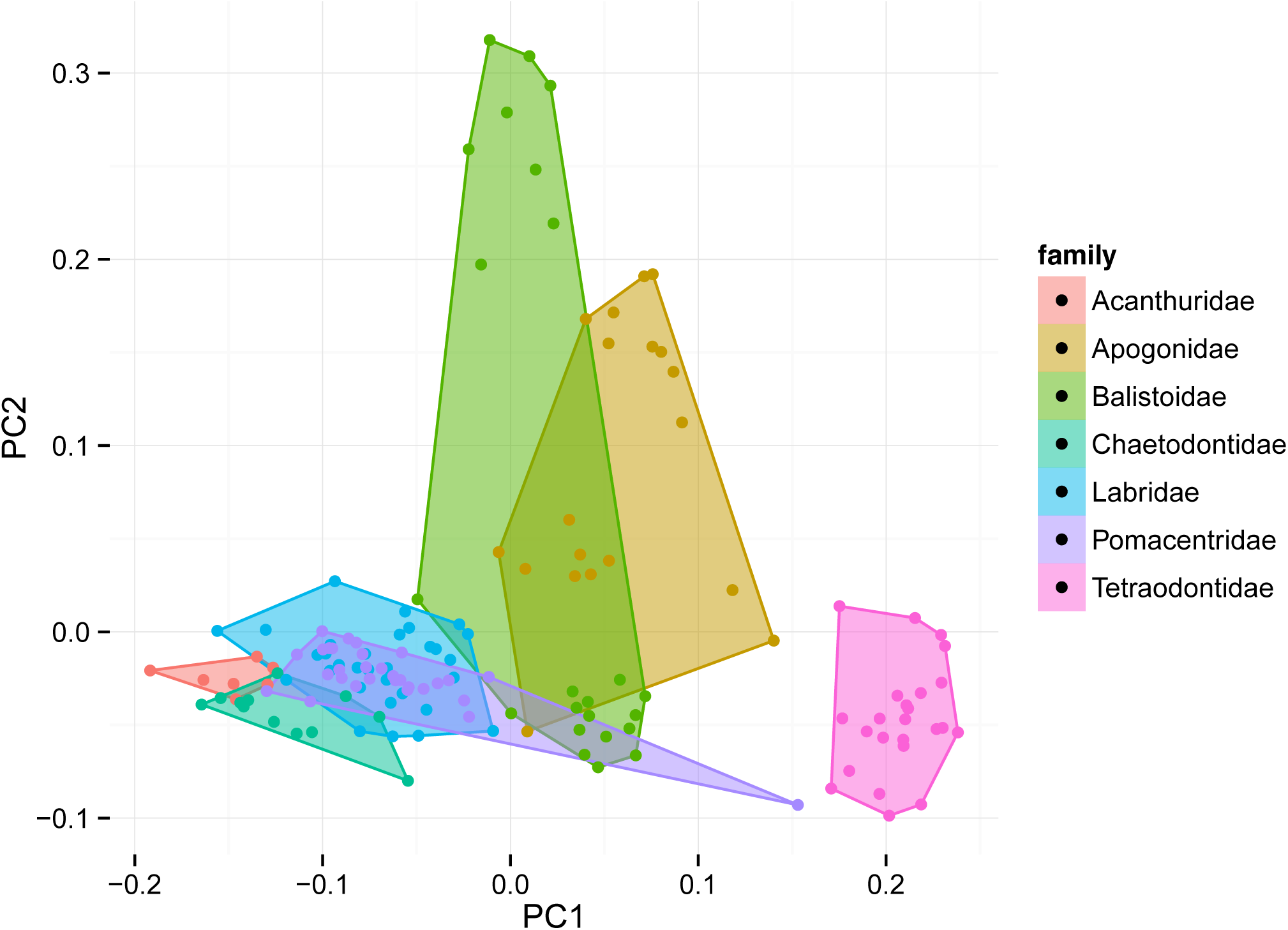
Morphospace for seven families of ray-finned fishes. Each point indicates a separate species; families are separated by colors. The convex hull for each family is drawn to show area of morphospace occupied by each family. Figures for other PC axes are present in the Supplemental Material.

**Figure 5:**
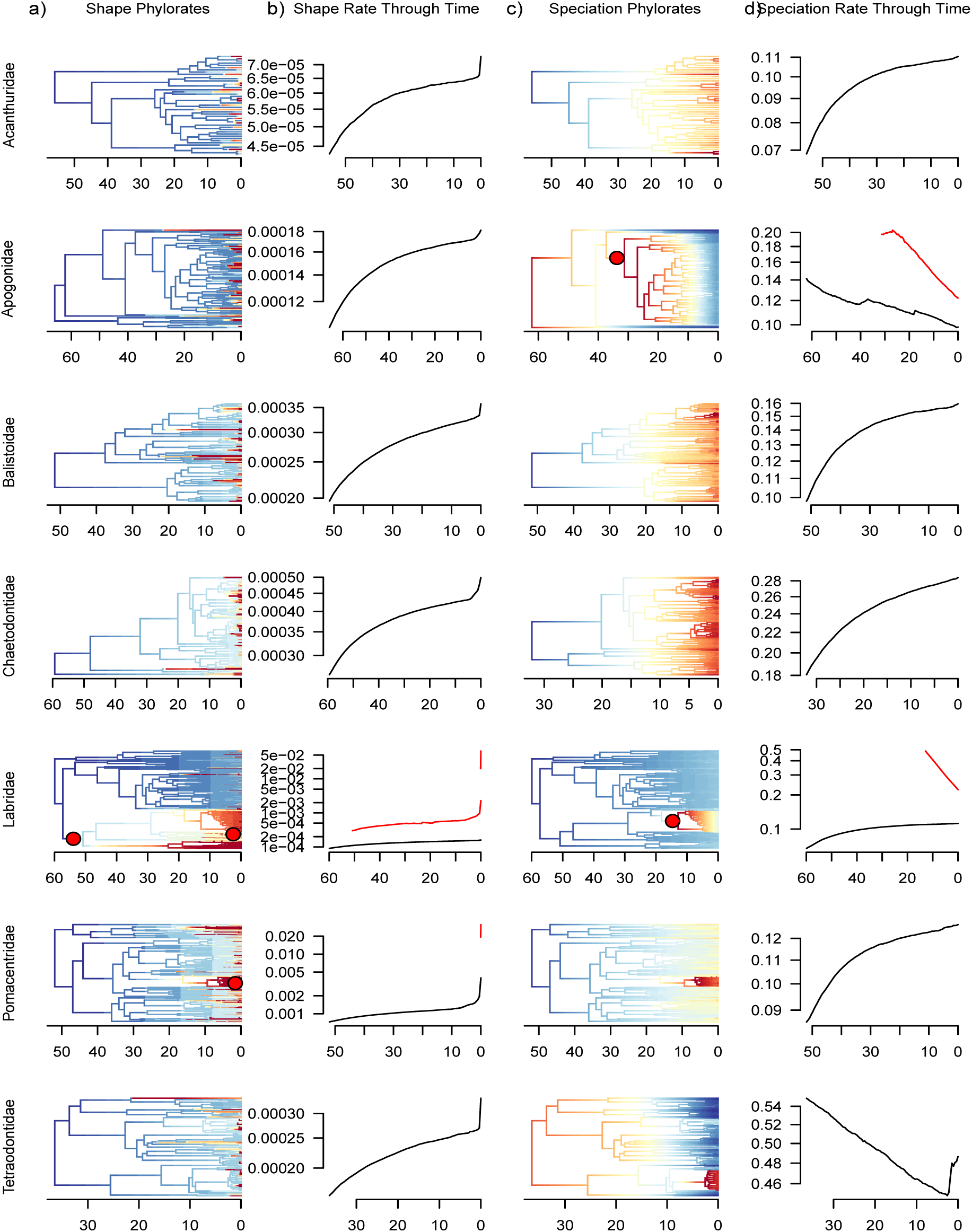
Rates of shape evolution for PC1 (a, b) and speciation (c, d) across seven families of fishes. Phylorate plots (a, c) color branch lengths by rates of shape evolution (a) and speciation (c), where warmer colors indicate faster rates of evolution. Significant rate shift events (pp > 0.95) are indicated on the phylorate plot as a red circle on the corresponding branch. Median log rates of shape evolution (b) and speciation (d) through time, where black lines indicate the background rate and red lines indicate the rate of evolution in a clade experiencing a significant shift in rate, corresponding to red circles in (a) or (c).

## Discussion

We have shown that crowdsourcing through Amazon Mechanical Turk is a tractable approach for generating reliable trait data at an unprecedented scale. Using this framework, it is possible to distribute thousands of images to workers, collect the data, and send it to a comparative analysis pipeline. We have also demonstrated that it is possible to identify the set of geometric morphometric landmarks that can be reliably captured by nonspecialists. We found that for certain landmarks there was significant between and within group disagreement. Based on median average deviation, points belonging to the opercular series and those that locating the distal margin of the dorsal and anal fins were particularly challenging, compared to the experts. Based on these results, nonspecialist turkers are unlikely to replace experts for all morphometric tasks. However, by digitizing less than 5% of our dataset with experts, we were able to identify groups of landmarks that exhibited extremely poor performance and excluded these. Furthermore, we were able to obtain biologically significant results from a dataset collected entirely by turkers. Through combining expert knowledge with the sheer scale of the Amazon Mechanical Turk workforce, it is possible to collect and assess large quantities of morphometric data, with an order of magnitude improvement in throughput over traditional approaches.

### Reliability

One advantage of the crowdsourced method we develop here is that inter-observer error can be readily assessed. Traditional geometric morphometric studies often rely on a single observer for practical reasons (the pool of trained geometric morphometricians is limited), and to avoid individually-driven systematic biases in data collection. Although this common practice may reduce bias, it also precludes meaningful assessment of differences among observers. Our results show that inter-observer variance can be substantial for some landmarks even among expert digitizers. Therefore, explicitly accounting for inter-observer error is critical to determine the efficacy of each individual landmark and the replicability of the study as a whole. Inter-observer error signals which landmarks can be relied on and which merit further consideration, as we have done in this analysis. The quantification of inter-observer error is a strict requirement of our workflow, as it would otherwise be impossible to arrive at a single consensus shape across several turkers working independently. This requirement ensures that inter-observer error is not ignored or bypassed due to the difficulty of assessing it.

In our analysis, we assessed the quality of a variety of landmarks between turkers and experts. Unsurprisingly, turkers performed exceptionally poorly for several landmarks requiring knowledge of fish anatomy. For example, the landmarks that describe the shape of the fish’s caudal fin asked workers to mark the distal tip of the first principal fin ray. Even when turkers are armed with a definition and a comparison between procurrent and principal fin rays, the experts’ experience and training allow them to substantially outperform turkers in identifying this point. Furthermore, experts generally had lower disagreement in their landmark placement when compared to turkers, even for landmarks that turkers found especially difficult. These differences between experts and MTurk workers have also been observed in image categorization tasks (Deng *et al.* 2009; Van Horn *et al.* 2015). However, it is possible that an improved training protocol could result in better collection of these difficult landmarks. Turkers have been found to perform well in extremely detailed video annotation tasks (Vondrick *et al.* 2013), provided that researchers conduct pre-task training and post-task validation. Implementing these pre-task requirements would be a straightforward avenue to improve accuracy for future work.

### The role of crowdsourced phenotypic data collection in modern comparative studies

The traditional way of collecting phenotypic data involves enormous researcher effort and significant morphological expertise. For example, Brusatte *et al.* (2014b) used a 853 character discrete character matrix for 150 taxa to estimate the rate of morphological evolution in the transition from theropod dinosaurs to modern birds. These data were collected over the course of 20 years as part of the Therapod Working Group (Brusatte *et al.* 2014a). O’Leary *et al.* (2013) combined the work of MorphoBank contributors (O’Leary & Kaufman 2011) with literature review to generate 4,541 characters for 86 species. Rabosky *et al.* (2013) examined 7,822 species of ray-finned fish and used a single quantitative measure (body size) collected from FishBase (Froese & Pauly 2014), whose data are contributed from the scientific literature by experts. All of these studies share the same requirement for intensive researcher effort, but the data collected is generally either broad (many species) or deep (many characters). In this study, we collected a phenotypically rich dataset across great taxonomic breadth. This approach can easily be scaled to permit unprecedented, massive comparative analyses on new, phenotypically rich datasets.

This method does not threaten to replace experienced morphologists. Though certain conspicuous landmarks can be rapidly collected by turkers, other types of analyses will require landmarks that can only be identified by experts and thus cannot use the high-throughput method presented here. Although this can likely be alleviated by implementing more sophisticated training regimes, the implicit anatomical knowledge that morphologists have must be made explicit in the form of a written protocol for turkers to follow. The cost of developing a clearer and simpler protocol that still captures the essence of the morphological characters of interest must be weighed against the benefit of higher-throughput from turker data collection, and for many such analyses this tradeoff is impractical. However, for such analyses where crowdsourcing is a viable alternative, our approach allows experts to move beyond data collection and into a role of developing training materials for nonspecialists and validating the data collected by crowdsourced workers.

Approaches involving statistical techniques like machine vision and natural language processing have yet to make significant headway in automatically collecting morphological data. Although methods to automatically measure leaves exist (Corney *et al.* 2012a; b), these require 2D specimens to eliminate parallax error, as well as high-contrast mounting paper backgrounds for effective automatic outline detection. More sophisticated methods for lower-quality images or organisms with more 3D structure have yet to be developed. Natural language processing of the scientific literature could potentially be used for automatic extraction of morphological characters using DeepDive (Peters *et al.* 2014; Shin *et al.* 2015), but it may require impractically large corpus sizes (Brill 2003; Halevy *et al.* 2009). Crowdsourcing can augment and enhance these statistical techniques. For example, the algorithm in Corney *et al.* (2012a) occasionally captures non-leaf objects and systematically underestimates leaf sizes. MTurk workers could improve this method by confirming the presence of a leaf in the image segment and measure the leaf size to ground truth the algorithm’s results.

A third alternative to using expert morphologists and crowdsourced workers to collect data is through citizen science. Citizen scientists are enthusiasts that volunteer to collect data or contribute annotations to a scientific endeavor. They can specialize in a particular field, such as birds, plants, or fungi. Compared to Amazon Mechanical Turk workers, citizen scientists are typically unpaid, but can produce higher quality work due to their expertise. For example, a study comparing citizen scientists and MTurk workers showed that for an image segmentation task MTurk workers had higher throughput and comparable accuracy to citizen scientists, but MTurk workers performed poorly when asked to identify birds to the species level (Van Horn *et al.* 2015).

### Suitability for other systems

Our novel pipeline to download images, upload them to Amazon MTurk, and process them using BAMM and BAMMtools showcases the ability to rapidly collect phenotypic data. Most of the time taken to collect these data were spent on waiting for worker results; however, a majority of the data had already been collected at the 1-hour mark. An online methodology could conceivably improve on this analysis time, by iteratively refining its results as new data streamed in from Amazon’s servers.

Although there are limitations in the type and accuracy of data that can be collected through MTurk crowdsourcing, even a simplified protocol can produce meaningful biological results that are concordant with previous hypotheses in these groups. We detected a significant shift in the rate of body shape evolution in Labridae, restricted to the wrasse tribes Labrini, Cheilini, and Scarini. The scarines and cheilines are mostly reef-associated (Froese & Pauly 2014), which has been proposed as an environment that drives diversification rate changes in marine teleosts (Alfaro *et al.* 2007; Cowman & Bellwood 2011; Price *et al.* 2011). These results suggest that evolution of body form may also be influenced by environmental association (Claverie & Wainwright 2014). Although the example we present here was necessarily limited, extending this technique to generate new phenotypic datasets for existing large phylogenetic trees such as fishes (Rabosky *et al.* 2013), birds (Jetz *et al.* 2012), mammals (Bininda-Emonds *et al.* 2007), and angiosperms (Zanne *et al.* 2014) would be straightforward, especially for taxa where image data are already aggregated in a database such as FishBase (Froese & Pauly 2014) or the Encyclopedia of Life (Parr *et al.* 2014).

Our approach hits a “sweet spot” on the three axes of expertise, effort, and computational complexity. We use researcher expertise to identify a comparative hypothesis, and design a data collection protocol to specifically test this hypothesis. Amazon Mechanical Turk supplies a large source of worker effort that collects data according to protocol. Finally, computational statistical techniques validate the accuracy of our data and identify outliers and other errors in data collection. Researchers do not have to spend time digitizing collections, workers need not generate biological hypotheses, and biologists will not have to solve open questions in the fields of machine vision and natural language processing in order to answer questions in comparative biology. The task of phenomic-scale data collection is split up and efficiently allocated according to the strengths of each role, without overly relying on any one axis to carry out the entire task.

Our work fills the niche of gathering phenotypic data across large radiations, which has been a challenging open research question (Burleigh *et al.* 2013). Even seemingly obvious phenotypes, such as the woodiness of plant species, are incomplete and sampled in a biased manner (FitzJohn *et al.* 2014), potentially misleading inference on a global scale. This method unlocks the potential of high-throughput data collection, and shifts the data bottleneck for morphological research onto acquiring suitable images for quantification, and developing higher-quality worker training regimens to enable collection of more sophisticated data. The burden is now on experienced taxonomists and morphologists to create protocols that are simple enough to be understood by MTurk workers, but comprehensive enough to test hypotheses of interest across the tree of life. Additionally, museums and other institutions must increase their efforts to make their biodiversity collections available digitally, including images suitable for morphological research. The problem of difficult-to-retrieve *dark data* is well-known (Heidorn 2008), but without either physical access to the collections or an image of the specimen, morphological data is impossible to acquire.

Our results suggest that, where possible, crowdsourcing should be an integral part of any large-scale morphological analysis. Crowdsourcing should play a key role in unlocking the “dark data” present in biodiversity collections by providing a high-throughput way to extract the phenotypic data present in specimens. Furthermore, coordinating efforts from digitizing museum collections, natural language processing and machine vision software, citizen scientists, expert morphologists and taxonomists, and crowdsourced Mechanical Turk workers would result in an extremely powerful pipeline that could generate a “phenoscape” across the tree of life.

## Acknowledgements

We thank XXX, YYY, and ZZZ for helpful comments on the manuscript, as well as T. Marcroft, B. Frederich, V. Liu, R. Aguilar, R. Ellingson, F. Pickens, C. LaRochelle, and the 22 Amazon Mechanical Turk workers that contributed their time and effort. We also thank D. Rabosky, B. Sidlauskas, M. McGee, A. Summers, and M. Burns for insightful discussions about fish morphology and digitization protocols. M. Venzon and T. Claverie provided unpublished figures that assisted this study. K. Staab and T. Kane allowed 156 undergraduate students to beta test the methods. This work was supported by an Encyclopedia of Life David M. Rubenstein Fellowship (EOL-33066-13), a Stephen and Ruth Wainwright Fellowship, and a UCLA Research and Conference Award to JC. Travel support to present this research was provided by the Society for Study of Evolution.

## Data Accessibility

All data are deposited online at the Encyclopedia of Life and Dryad.

## Author contributions

Conceived and designed the experiments: JC MEA. Performed the experiments: JC. Analyzed the data: JC. Contributed reagents/materials/analysis tools: JC MEA. Wrote the paper: JC MEA.

